# Development and Validation of a Novel Instrument to Capture Companion Dog Mortality Data: The Dog Aging Project End of Life Survey

**DOI:** 10.1101/2023.04.01.535178

**Authors:** Kellyn E. McNulty, Kate E. Creevy, Annette Fitzpatrick, Vanessa Wilkins, Brian G. Barnett, DAP Consortium, Audrey Ruple

## Abstract

**Objective:** The researchers and clinicians within the Dog Aging Project (DAP), a longitudinal cohort study of aging in companion dogs, created and validated a novel survey instrument titled the End of Life Survey (EOLS) to gather owner-reported mortality data about companion dogs.

**Sample:** Bereaved dog owners who participated in the refinement, face validity assessment, or reliability assessment of the EOLS (n=42) and/or completed the entire survey between January 20 and March 24, 2021 (n=646).

**Procedures:** The EOLS was created and modified by veterinary health professionals and human gerontology experts using published literature, clinical veterinary experience, previously created DAP surveys, and feedback from a pilot study conducted with bereaved dog owners. The EOLS was subjected to qualitative validation methods and post-hoc free-text analysis to evaluate its ability to thoroughly capture scientifically relevant aspects of companion dogs’ death.

**Results:** The EOLS was well-received with excellent face validity as assessed by dog owners and experts. The EOLS had fair to substantial reliability for the three validation themes: cause of death (kappa = 0.73; 95% CI [0.5-0.95]), perimortem quality of life (kappa = 0.49; 95% CI [0.26-0.73]), and reason for euthanasia (kappa = 0.3; 95% CI [0.08-0.52]) and had no need for any substantial content alterations based on free-text analysis.

**Clinical Relevance:** The EOLS has proven to be a well-accepted, comprehensive, and valid instrument for capturing owner-reported companion dog mortality data and has the potential to enhance veterinarians’ ability to care for the aging dog population by illuminating their understanding of companion dogs’ end-of-life experiences.

## Introduction

A common goal among aging studies is elucidating the causes and consequences of aging in order to improve lifespan and/or healthspan, the period of one’s lifespan spent free from chronic or debilitating conditions. As such, the primary endpoint of aging studies is death. This is recorded on human death certificates as both the specific cause of death, defined as the underlying medical condition, disease, or injury that results in death, as well as the manner of death, defined as the way in which a death occurs (e.g. accident, homicide etc.). Given the popularity and intrinsic value of companion dogs in developed countries and the myriad of similarities between canine and human physiology, environmental exposures, and lifestyle, aging research investigating the cause and manner of death in companion dogs presents a significant opportunity to benefit the well-being of both species.^1^

Before factors that influence healthy lifespan can be discovered, aging studies must first obtain the endpoints necessary to make meaningful conclusions. To accurately capture this mortality data in humans, studies must contend with under-reported or incorrectly reported cause of death and/or manner of death information when individuals die at home,^2, 3^ when death certificates are improperly completed,^4, 5^ when individuals have several comorbidities at the time of death, and when autopsies are not performed.^6^ Veterinary studies investigating the cause and manner of death in companion dogs face similar obstacles as well as additional unique challenges.

In the United States, there is no widespread standardized system for reporting or coding companion dogs’ cause or manner of death. The few prominent databases commonly used for large retrospective companion dog studies are not thought to be representative of the national population of companion dogs (e.g. insured dogs or dogs seen at corporate or teaching veterinary hospitals).^7–10^ Additionally, because some companion dogs die at home, veterinarians may not be informed of their patients’ deaths, and thus have no opportunity to capture information about the causes or manners of their deaths. Therefore, dog owners represent a key source of data on companion dog death with the potential to overcome these obstacles.

Another challenge encountered when analyzing end of life circumstances in companion dogs is that euthanasia is a common manner of death in this species.^7, 8, 11–13^ In the United States, there are no legal restrictions regarding the reasoning for pursuing euthanasia. Therefore, euthanasia may be performed for many reasons (poor quality of life, aggressive behavior, financial hardship etc.)^14–17^ and theoretically can occur at any point during a dog’s lifetime. Additionally, owners whose dogs have the same cause of death (e.g., metastatic cancer) may choose euthanasia for different reasons (e.g., poor quality of life or poor prognosis), while owners whose dogs have different causes of death may choose euthanasia for the same reason (e.g., pain and suffering). Consequently, any attempt to comprehensively capture information concerning companion dog death must ask the owner about the dog’s cause of death, manner of death (i.e. euthanasia vs. unassisted death) and any factors that may have influenced their decision to euthanize.

The investigators within the Dog Aging Project (DAP),^18^ a prospective, longitudinal cohort study of aging in companion dogs in the United States, have developed a strategy to address this complex challenge. Similar to human longitudinal aging studies, which utilize death certificates, medical records, physician panel review, next-of-kin or proxy interviews, and autopsies,^19–21^ the DAP team has planned a tiered data-collection system for recording cause of death, manner of death, and circumstances surrounding the death of participating dogs. The first tier of this data-collection system required the development of a novel survey instrument titled the End of Life Survey (EOLS), which was designed to be completed by dog owners.

Previously published questionnaires created to capture companion dog cause of death are often selective in the breeds evaluated,^12, 22, 23^ fail to investigate the reason(s) respondents report “old age” as a dog’s cause of death,^12, 23–25^ and/or only explore owners’ motivations for euthanasia with regard to specific, as opposed to all, causes of death.^26–28^ As such, the DAP team developed the EOLS to acquire more detailed and comprehensive information about participant dogs’ deaths directly from owners. The aims of this study were to describe the development of the EOLS and to demonstrate its validity as a data-collection tool for capturing the most salient factors necessary to understand companion dog death, namely owner-reported cause of death, reason for euthanasia, and perimortem quality of life.

## Materials and Methods

### The DAP Overview

The DAP has been previously described.^18^ Briefly, the DAP interacts with participants primarily through personalized online portals, and data are collected and managed using REDCap (Research Electronic Data Capture) tools hosted at the University of Washington.^29, 30^ All owners complete an initial comprehensive survey, called the Health and Life Experience Survey (HLES), designed to obtain information about the dog’s husbandry, physical and social environment, and health. Data collected by this survey are updated annually.

The DAP is informed of a participating dog’s death through one of several mechanisms. These include participants self-reporting whether the dog remains a member of their household prior to completing research tasks, participants calling or emailing the study team, or participants updating the “major event form,” a communication tool within their online portal. Once an owner reports the death of their participating dog through any of these mechanisms, they are invited to complete the EOLS.

### The EOLS Overview

The EOLS was created and validated by a veterinary resident with prior general practice experience (KEM), a veterinary internist (KEC), a veterinary epidemiologist (AR), and a human health epidemiologist (AF). This was accomplished through reviewing relevant literature, consulting experts in end-of-life data collection in human populations, conducting a pilot study, and performing a qualitative validation assessment. The EOLS items were created to gather detailed information related to companion dog death **(Table 1)**. Among these items, three (cause of death, reason for euthanasia, and perimortem quality of life) were chosen by the authors to be validation themes as they were the most essential for understanding end of life circumstances in companion dogs and were also the least objective.

**Table 1.** Questions and answer types contained within the Dog Aging Project (DAP) End of Life Survey (EOLS).

| Question | Answer Type |
| --- | --- |
| **Q1** Was your dog diagnosed with health conditions in any of the following categories [since the last time we collected health history information from your dog]? | C+OPD |
| **Q2** Please indicate the date of your dog’s death. If you know the month but not the exact day, do your best to estimate or simply select the 15th of the month. | automated calendar date |
| **Q3** Thinking back on the last two weeks of your dog’s life, did they exhibit any of the following characteristics related to the aging process, or general old age? | C+OPD |
| **Q4** Thinking back on the last two weeks of your dog’s life, did they exhibit any of the following medical symptoms or signs? | C+OPD |
| **\*Q5** In the two weeks before your dog’s death, how would you describe their quality of life? | LT |
| **Q6** At what point in your dog's lifetime do you think that their quality of life first began to decline? | C |
| **Q6a** If you noticed a decline in your dog's quality of life, what contributed most significantly to this decline? | C+OPD |
| **Q7** Was your dog's health evaluated by or discussed with (either in person or by phone) a veterinarian within 2 weeks of the date of their death (including the day of death)? | Y/N/U |
| **Q7a** Did your dog stay at a veterinary clinic or hospital to receive any type of treatment in the 2 weeks prior to their date of death? | Y/N/U |
| **Q7a-1** How long did your dog stay in the veterinary clinic or hospital? | C/U |
| **Q7b** Did your dog undergo sedation, anesthesia, or surgery for any reason within 2 weeks of their date of death? | Y/N/U |
| **Q7c** Given the treatment option you chose (including no treatment), what was your understanding of your dog's prognosis (probability of cure or successful long-term management)? | LT/U |
| **Q8** Indicate the location of your dog's death. | C+OPD/U |
| **Q9** Was anyone present at your dog's death? | Y/N/U |
| **Q9a** Who was present at your dog's death? | C+OPD |
| **Q10** Was your dog euthanized (put to sleep or put down)? | Y/N |
| **Q10a** Who performed the euthanasia? | C+OPD |
| **\*Q10b** Which of the following was the most significant contributing factor in your decision to choose euthanasia? | C+OPD |
| **Q10c** Which additional factors contributed in any way (large or small) to your decision to choose euthanasia? | C+OPD |
| **\*Q11** Which one of the following categories best describes your dog's cause of death (whether or not euthanasia was performed)? | C+OPD |
| **Q12** Which of the following categories best describes the second most important contributor to your dog's cause of death (whether or not euthanasia was performed)? | C+OPD |
| <u>If Q11 Cause of Death = OLD AGE</u><br>**OLD AGE (a)** Which of these characteristics of old age was the <u>most significant</u> cause of your dog's death? | C+OPD |
| <u>If Q11 Cause of Death = ILLNESS</u><br>**ILLNESS (a)** What type of fatal illness was the cause of your dog's death? | C/U |
| <u>If Q11 Cause of Death = ILLNESS and ILLNESS (a) = CANCER</u><br>**CANCER (a)** If possible, please indicate which body system(s) was involved in your dog's cancer. | C+OPD/U |
| **CANCER (b):** Do you know the name of the specific cancer that led to your dog's death? | Y/N |
| Please provide the name of the specific cancer | FT |
| <u>If Q11 Cause of Death = ILLNESS and ILLNESS (a) = INFECTION</u><br>**INFECTION (a)** If possible, please indicate which infection your dog had. | C+OPD/U |
| **INFECTION (b)** If possible, please indicate which body system was involved in the infection. | C+OPD/U |
| **If Q11 Cause of Death = ILLNESS and ILLNESS (a) = OTHER-ILLNESS**<br>**OTHER-ILLNESS (a)** If possible, please indicate which body system was involved in your dog's illness. | C+OPD/U |
| **OTHER-ILLNESS (b)** Do you know the specific medical diagnosis that led to your dog's death? | Y/N |
| Please provide the name of the specific diagnosis | FT |
| **If Q11 Cause of Death = ILLNESS** |  |
| **ILLNESS (b)** At what point in your dog's lifetime did you first become aware of the illness that would ultimately be the cause of their death? | C |
| **ILLNESS (c)** Was your dog being treated for this illness at the time of their death? | C+OPD |
| **If Q11 Cause of Death = TRAUMA**<br>**TRAUMA (a)** What type of trauma was the cause of your dog's death? | C+OPD/U |
| **If Q11 Cause of Death = TOXIN**<br>**TOXIN (a)** What type of toxin was the cause of your dog's death? | C+OPD/U |
| **If Q12 = Old Age**<br>**Old Age COD #2** Which of these characteristics of old age was the most significant reason you listed old age as the second most important contributor to your dog's cause of death? | C+OPD |
| **Q13** If there is anything else that you would like us to know about the circumstances surrounding your dog's death, please share it here: | FT |

Y/N = yes or no, U = unknown or unsure, LT = Likert-type, C = categorical (as the sole response option), FT = free-text (as the sole response option), C+OPD = categorical + free-text (as "other, please describe"), \* = validation themes analyzed with Cohen's kappa coefficient

### The EOLS Creation

Clinical veterinary experience informed the development of most survey items. Items related to perimortem quality of life and euthanasia were created based on published reports of owners’ perception of their dog’s quality of life^14, 31, 32^ and the reason(s) owners pursue euthanasia.^14–17^ Cause of death items were constructed based on mortality studies in dogs^8, 9, 12, 13, 33^ and previously published but unvalidated companion dog cause of death surveys.^12, 22–26^ Previous reports on physical^34–37^ and behavioral^38–46^ changes associated with companion dog aging were utilized to develop the items related to “old age” characteristics. When applicable, items were constructed to align with terminology used in similar components of the HLES to facilitate data mapping between instruments.

The EOLS contains eight categorical causes of death **(Table 2)**. For some causes of death, additional items were nested within responses so that a specific medical diagnosis could be captured, if known. For example: “illness/disease” => “cancer” => “spleen” => free-text entry “hemangiosarcoma.” The EOLS contains eight categorical reasons for euthanasia (**Table 3**), and a 7-point Likert-type scale is used to assess quality of life (**Table 4**). Various questions and answer types were used in the EOLS (Table 1) including an optional free-text item at the conclusion of the survey allowing participants to share additional narrative information.

**Table 2.** Primary categorical cause of death in 646 dogs as reported by owners and captured in the Dog Aging Project (DAP) End of Life Survey (EOLS)

| **Cause of death** | **Number of dogs** |
| --- | --- |
| Illness or disease | n = 382 (59.1%) |
| Old age | n = 190 (29.4%) |
| Other | n = 24 (3.7%) |
| Trauma or injury | n = 21 (3.3%) |
| Sudden death | n = 18 (2.8%) |
| Sedation, anesthesia or surgery | n = 6 (0.9%) |
| Toxin | n = 3 (0.5%) |
| Behavior problem | n = 1 (0.2%) |
| Personal factors | n = 1 (0.2%) |

**Table 3.** Primary reason for euthanasia in 536 dogs as reported by owners and captured in the Dog Aging Project (DAP) End of Life Survey (EOLS)

| Reason for euthanasia | Number of dogs |
| --- | --- |
| Pain and/or suffering | n = 260 (48.5%) |
| Poor quality of life | n = 133 (24.8%) |
| Poor prognosis | n = 105 (19.6%) |
| Unmanageable medical problem | n = 26 (4.9%) |
| Other | n = 7 (1.3%) |
| Unmanageable behavior problem | n = 2 (0.4%) |
| Fear of harm to another animal or person | n = 2 (0.4%) |
| Incompatible with home situation | n = 1 (0.2%) |
| Cost of care | n = 0 (0%) |

**Table 4.** Quality of life during the two weeks prior to death in 646 dogs as reported by owners and captured in the Dog Aging Project (DAP) End of Life Survey (EOLS)

| Quality of life | Number of dogs |
| --- | --- |
| Always bad days | n = 10 (1.5%) |
| Almost entirely bad days | n = 57 (8.8%) |
| More bad days than good days | n = 152 (23.5%) |
| Equal number of good days and bad days | n = 89 (13.8%) |
| More good days than bad days | n = 145 (22.4%) |
| Almost entirely good days | n = 131 (20.3%) |
| Always good days | n = 62 (9.6%) |

### The EOLS Refinement and Face Validity Assessment

A total of 42 dog owners participated in the refinement, face validity assessment, and reliability assessment of the EOLS. A preliminary version of the EOLS was presented to two recently bereaved dog owners and modified for clarity based upon their feedback. A pilot study was performed with 13 participants from two convenience sample groups: three members of the DAP community advisory board (CAB)^18^ whose dogs were not study participants and ten DAP participants whose participating dog had died within the prior month.

All pilot participants completed the EOLS and subsequently provided feedback by sharing additional narrative information at the end of the survey (CAB participants=1; DAP participants=10), responding to a set of emailed questions (CAB participants=3; DAP participants=6), and/or participating in a structured phone interview with one of the authors (KEM) (CAB participants=0; DAP participants=5). The emailed questions prompted participants to comment on survey timing, technical difficulties, and comprehensiveness and clarity of the items. Each interviewed owner was invited to describe their dog’s cause of death and reason(s) for euthanasia, and to provide any additional thoughts regarding the survey.

Feedback from this pilot study was used to assess the functionality, clarity, comprehensiveness, and qualitative face validity of the instrument. Qualitative face validity was further assessed by discussing each item with a panel of experts including the authors, a palliative care veterinarian (LM), a human subjects research regulatory specialist (ECJ), and a communications expert within the DAP (AK). All feedback was used to revise and produce the final version of the EOLS.^47^

### The EOLS Reliability Assessment

One month prior to the EOLS launch, a convenience sample of DAP participants whose dogs had been reported as deceased within the prior month were invited to be part of a qualitative reliability assessment that compared participants’ verbal description of their dogs’ death to their subsequent EOLS responses. Participants were asked to schedule an interview through an online scheduling program (Calendly; Calendly LLC), attend a phone or virtual interview with one of the authors (KEM), and then complete the EOLS in their online portal within the month following the interview. Twenty-eight of the forty invited participants scheduled and completed an interview, and twenty-seven of those completed the EOLS within the following month. Thus, there were 27 participants involved in the qualitative reliability assessment.

After obtaining participant consent, each phone (Rev Call Recorder; Rev.com Inc) or video (Zoom; Zoom Video Communications Inc) session was recorded. Each participant was asked to provide a detailed description of the events surrounding their dog’s death. If the validation themes of the EOLS (cause of death, reason for euthanasia, perimortem quality of life) were not discussed in their spontaneous description, the interviewer used scripted open-ended questions to obtain that information. The responses collected in this proxy interview (PI) were used by the interviewer to complete the relevant sections of the EOLS immediately following the interview, which generated a proxy interview EOLS (PI-EOLS) for each participant.

A second veterinarian (BGB) reviewed the recordings for half of the interviews and completed a second version of the PI-EOLS to increase confidence in the interviewer’s interpretation of participant responses. The two PI-EOLS versions were compared, and there were no major discrepancies. Minor discrepancies were reconciled through discussion and the final consensus PI-EOLS responses were then compared to each participant’s subsequent EOLS responses.

### The EOLS Launch

The EOLS^47^ was integrated into the DAP online platform on January 20, 2021. On that date, owners of all DAP participant dogs previously reported as deceased were invited to complete the EOLS. Since that date, any subsequently reported death of a DAP participant dog generates an invitation for the owner to complete the EOLS. Participants are given 40 days to complete the EOLS. All EOLS survey responses submitted between January 20 and March 24, 2021 from participants who had enrolled in the DAP and completed the HLES by December 31, 2020 were analyzed in this study.

### The EOLS Free-Text Assessment

Two types of free-text responses were analyzed in this study. The optional item at the end of the EOLS records any additional free-text narrative information participants wish to share. These free-text responses were read in their entirety and an inductive approach was used to code the qualitative data using the comments provided by respondents to develop the themes.^48^ These data were reviewed to identify any broad themes relevant to understanding companion dog death that were not already addressed in the EOLS.

The EOLS contains 20 items in which the response “other, please describe” and an associated free-text box is offered among the list of categorical responses. These free-text entries were read and coded by being placed into one of the following four categories: 1) duplicate of an available response choice for that item, 2) duplicate of an available response choice for a different item, 3) novel information relevant to EOLS objectives, and 4) novel information not relevant to EOLS objectives. These coded responses were analyzed to indirectly assess clarity and comprehensiveness of provided response choices and to identify frequently reported, relevant response choices not offered in the EOLS.

Novel, relevant themes described in the additional narrative information and free-text responses from “other, please describe” items which were repeated by >5 individuals and represented >1% of the response rate for that item were evaluated in more detail.

### Statistical Analysis

For the reliability assessment, agreement between participants’ PI-EOLS and EOLS responses to Q5, Q10b, Q11 (Table 1) was calculated using Cohen’s kappa coefficient. Kappa scores and 95% confidence intervals were reported for each EOLS theme evaluated. Kappa values were interpreted as follows: <0 = no agreement, 0.0-0.2 = slight agreement, 0.21-0.4 = fair agreement, 0.41-0.6 = moderate agreement, 0.61-0.8 = substantial agreement, 0.81-1.0 = almost perfect agreement.^49^

For the additional narrative free-text assessment, the identified overarching themes were assessed as counts and percentages within the context of all participants who responded to this optional item. For the “other, please describe” free-text assessment, the four coded categories were assessed as counts and percentages within the context of the total number of participants who responded to each item, including those who selected an available response choice.

## Results

### The EOLS Refinement and Face Validity Assessment

Edits and alterations were made to the EOLS in response to pilot and expert panel feedback. Item phrasing, order, and survey logic were also modified to enhance clarity and ease of completion. The expert panel concluded that the EOLS content and format were appropriate to capture scientifically relevant information regarding companion dog death in a suitably sensitive manner. Functionality and clarity of the EOLS were both reported to be excellent by pilot testers; none of the thirteen pilot testers experienced technical difficulties, found questions unclear or confusing, or found the time for completion to be burdensome. The median completion time for those pilot testers who recorded their time (n=9) was 10 minutes with a range of 5-30 minutes.

Eleven of thirteen (84.6%) pilot participants’ survey feedback was positive and indicated that the EOLS was comprehensive, straightforward, and compassionate. Quotes from several participants included: “I felt that this survey addressed everything I wanted to share about the passing of my dog.” “The questions were very appropriate…easy to read and easy to understand.” “The final paragraph [at the close of the survey] is super. It demonstrates to me that the study team really cares about dogs and their owners.” “[Completing this survey] fulfilled a desire to give [my dog’s] life more meaning…and it felt good doing it…It felt healing.” While the remaining two pilot participants initially reported that the EOLS items did not capture everything they wanted to share about their dog’s passing, both participants said that the ability to provide additional narrative information at the end of the survey alleviated their initial concerns and enabled them to tell their dog’s story the way they desired. Furthermore, for both participants, their dogs’ cause(s) of death, reason(s) for euthanasia, and perimortem quality of life, as revealed in the interview, were all accurately captured by EOLS items.

### The EOLS Reliability Assessment

Twenty-seven paired EOLS and PI-EOLS responses were generated in the reliability assessment to evaluate the validation themes cause of death and perimortem quality of life. Four of these dogs died unassisted, so only 23 paired responses were available for assessment of the validation theme reason for euthanasia.

Agreement between EOLS and PI-EOLS responses was substantial for cause of death (kappa = 0.73; 95% CI [0.5-0.95]). Among the 27 paired EOLS and PI-EOLS responses, 17 (63%) had complete agreement on the cause of death to the deepest level of nested questions. An additional five (18.5%) had complete agreement on the cause of death, but EOLS and PI-EOLS responses followed different nested pathways to the final diagnosis (e.g. “illness/disease” => “organ system” => “endocrine system” => free-text “Cushing’s disease,” vs. “illness/disease” => “cancer” => “pituitary gland” => free-text “Cushing’s disease”). In two interviews (7.4%), the participant identified two causes of death but failed to definitively commit to one being primary vs secondary, which led to EOLS and PI-EOLS responses containing the same causes of death but different identification of primary vs. secondary. In three (11.1%) cases, participants gave categorically different responses in the recorded interview and subsequent EOLS instrument.

Agreement between EOLS and PI-EOLS responses was moderate for perimortem quality of life (kappa = 0.49; 95% CI [0.26-0.73]). Among the 27 paired EOLS and PI-EOLS responses, 16 (59.3%) had complete agreement on perimortem quality of life as captured by a 7-point Likert-type scale. Six (22.2%) paired responses differed by one degree on the Likert-type scale; four (14.8%) paired responses differed by two degrees on the Likert-type scale. A single pair of responses differed by four degrees on the Likert-type scale.

Agreement between EOLS and PI-EOLS responses was fair for reason for euthanasia (kappa = 0.3; 95% CI [0.08-0.52]). Respondents who euthanized their dog could indicate multiple reasons for euthanasia with one being designated as the primary reason. Among the 23 reliability assessment respondents who euthanized their dog, a median of three (range 1-4) reasons for euthanasia were indicated in both EOLS and PI-EOLS responses. Considering all reasons for euthanasia reported in the paired EOLS and PI-EOLS responses, seven (30.4%) displayed complete agreement on every reason for euthanasia, and nine (39.1%) displayed complete agreement except for a single reason for euthanasia which was present in EOLS or PI-EOLS responses and omitted from the other. Considering only the reason designated as the primary reason for euthanasia, EOLS and PI-EOLS responses were concordant in 8 out of 23 (35%). However, for almost all of the paired EOLS and PI-EOLS responses, the reason for euthanasia designated as primary in one was reported as one of the overall reasons for euthanasia in the other.

### The EOLS Responses

A total of 793 DAP participants reported the death of their dog between project enrollment on December 26, 2019 and March 24, 2021. All were invited to complete the EOLS and given 40 days to do so. As of March 24, there were 29 participants who had not yet completed the EOLS, but for whom 40 days had not elapsed; these participants were excluded from analysis. Of the 764 remaining participants, 655 completed the survey, 8 started the survey but did not complete it within the 40-day window, 10 opened the survey but chose to opt out, and 91 never opened the survey for an overall response rate of 85.7%. Nine participants who completed the EOLS had not completed HLES prior to December 31, 2020 and were excluded from analysis. This produced a total of 646 EOLS responses with matching complete HLES data for analysis. Of these 646 respondents, 536 (83.0%) euthanized their dog. The primary cause of death, primary reason for euthanasia, and perimortem quality of life assessment for this cohort are presented in Tables 2, 3, and 4, respectively.

### The EOLS Free-Text Assessment: General

The EOLS contained an optional free-text item at the conclusion of the survey and 20 items with the free-text response choice “other, please describe.” All participants saw the concluding free-text item; however, due to survey logic, not all participants saw every “other, please describe” item. Among 646 EOLS respondents, 519 (80.3%) utilized one or both of the analyzed free-text options; 266 respondents (41.2%) utilized the “other, please describe” free-text option at least once, and 467 respondents (72.3%) provided additional free-text narrative information at the end of the survey. A median of 13.5 (range 0-83) participants chose “other, please describe” for each of the 20 “other, please describe” items. For almost half (45%) of these items, fewer than 10 participants per item selected the response choice “other, please describe.”

Five of seven themes identified in the additional narrative section at the conclusion of the EOLS were addressed elsewhere in the survey **(Table 5)**. The two most prominent additional narrative themes focused on the timeline of events surrounding the dog’s death and the owner’s thoughts, emotions, and experiences related to their dog’s death, which were not among the objectives of the EOLS.

**Table 5.** Themes identified within 467 participants’ free-text additional narrative information at the conclusion of the Dog Aging Project (DAP) End of Life Survey (EOLS)

| Theme | Number of responses containing each theme |
| --- | --- |
| Depiction of the owner’s own experience of the dog’s death | n = 437 (93.6%) |
| Detailed chronology of events surrounding the dog’s death | n = 399 (85.4%) |
| More detailed explanation of the dog’s medical symptoms | n = 306 (65.5%) |
| More detailed explanation of the dog’s quality of life and “old age” characteristics | n = 297 (63.6%) |
| More detailed explanation of the dog’s comorbidities | n = 256 (54.8%) |
| More detailed explanation of the treatments pursued at the end of the dog’s life | n = 115 (24.6%) |
| More detailed explanation of the diagnostics performed at the end of the dog’s life | n = 51 (10.9%) |

Ninety-five percent of the 7,799 participant responses to the “other, please describe” items selected one of the provided response choices leaving only 389 responses for investigation of the free-text responses associated with these items **(Figure 1)**. Half (n=196, 50.4%) of the free-text responses restated a response choice already provided for the given item. Fifty-five (14.1%) free-text responses for a given item restated a response choice that was already provided for a different item. This category was predominantly (n=40, 72.7%) composed of participants who used “other, please describe” in the item pertaining to “old age” characteristics to provide free-text entries that matched response choices already provided in the item pertaining to medical symptoms or vice versa. Additionally, when specifying cause of death under the category of “old age”, half (n=6, 50%) of participants’ free-text entries were response choices provided under a different categorical cause of death. Participants rarely (n=23, 5.9%) provided irrelevant free-text entries (e.g. “he just wasn’t well”). About one third of free-text entries (n=115, 29.6%) represented novel, relevant information; however, most of these responses were unique to that participant and not repeated.

**Figure 1.**
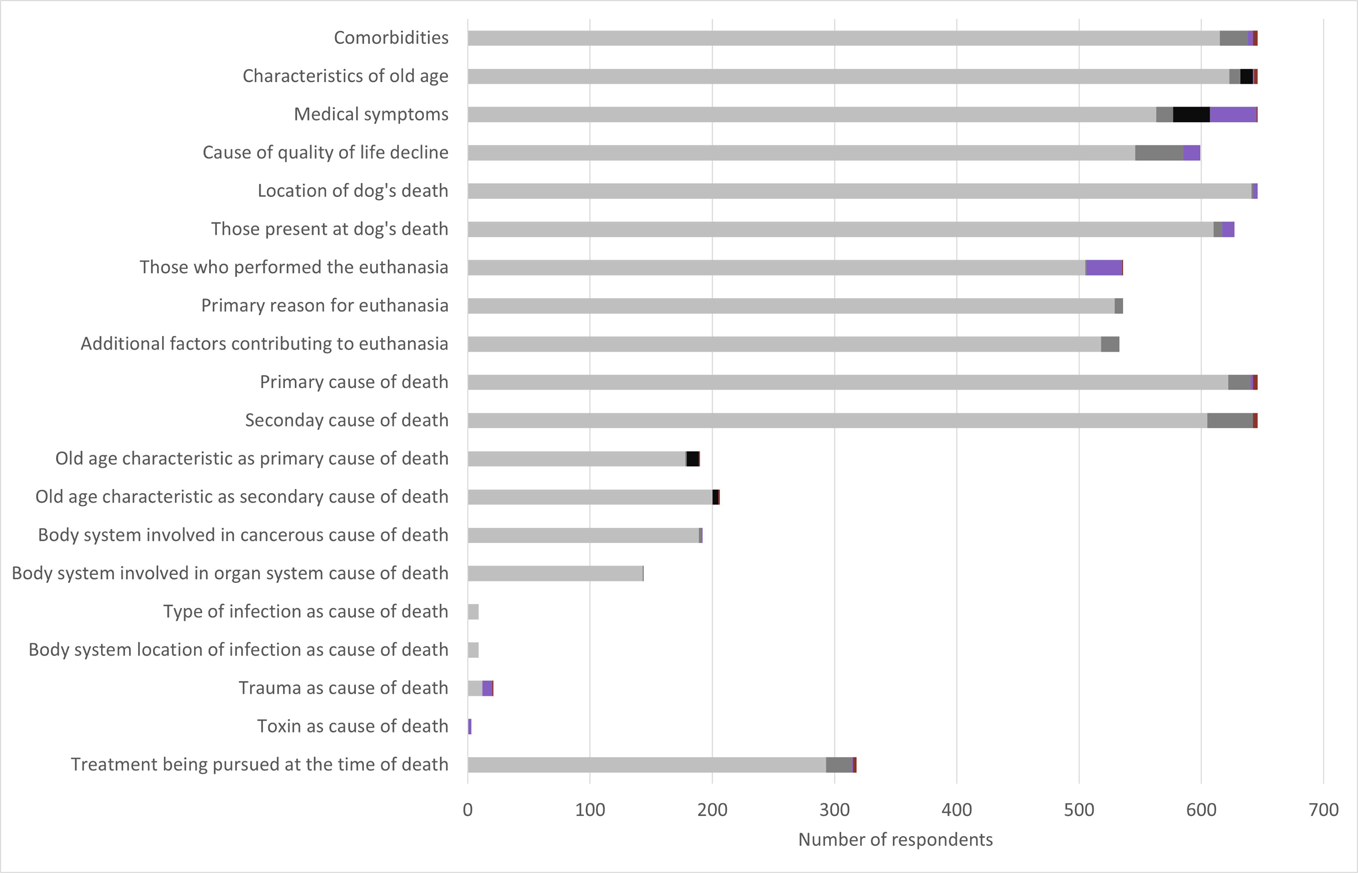
Twenty items within the Dog Aging Project (DAP) End of Life Survey (EOLS) offer a list of categorical response options including the option “other, please describe”. Among 646 survey respondents, the bar chart shows the number of respondents for each item who chose one of the provided response choices (light grey) or chose “other, please describe” and provided a free-text response. The “other, please describe” free-text responses were read and coded into the following categories: duplicate of an available response choice for that item (dark grey), duplicate of an available response choice for a different item (black), novel information relevant to EOLS objectives (purple), and novel information not relevant to EOLS objectives (red).

There were only two “other, please describe” items that contained free-text responses which were repeated by >5 respondents and represented >1% of the response rate for the given item. One of these was the item asking who performed their dog’s euthanasia; twenty-nine (5.4%) participants reported in the free-text that their dog was euthanized by an at-home veterinary euthanasia or hospice service. The second of these was the item asking who was present at their dog’s death; ten (1.5%) respondents reported in the free-text that another dog was present.

### The EOLS Free-Text Assessment Specifics: Cause of death and reason for euthanasia

Only twenty-four out of 646 (3.7%) participants utilized the option of “other, please describe” when reporting their dog’s primary cause of death. Of these, a majority (n=18, 75.0%) restated a provided response choice. Four free-text responses did not specify a scientifically relevant cause of death, and two provided relevant information that was not repeated. The option of “other, please describe” was only utilized by seven out of the 536 (1.3%) participants who had euthanized their dog to describe their dog’s primary reason for euthanasia, and all seven restated one of the provided response choices.

## Discussion

This manuscript describes the development and validation of the EOLS, a novel survey instrument designed by the DAP to capture scientifically important factors related to companion dog death. The EOLS was shown to be widely accepted by owners, comprehensive in its data collection, and both valid and reliable. Face validity was assessed by a panel of experts who confirmed that the EOLS successfully captured all scientifically relevant information pertaining to companion dogs’ death and by pilot study participants who reported the EOLS to be thorough, clear, compassionate, and not burdensome to complete. The investigative team utilized a qualitative reliability assessment to compare participants’ verbal descriptions of their dogs’ deaths to their digitally completed EOLS responses. This analysis revealed substantial, moderate, and fair agreement for EOLS validation themes: cause of death, perimortem quality of life, and reason for euthanasia, respectively.

There are several potential explanations for the fair Kappa agreement between PI-EOLS and EOLS responses for reason for euthanasia. While euthanasia decisions are almost invariably multifactorial,^14–17^ it was necessary to ask EOLS respondents to prioritize one reason as the primary reason to generate an analyzable data structure. If several reasons for euthanasia were equally important, participants may have selected a different primary reason when completing the EOLS than during the interview. Additionally, to avoid narrowing participants’ thoughts on this subject, the interviewer asked open-ended questions regarding overall reasons for euthanasia and recorded the reason mentioned first as the primary reason. More closed-ended interview questions and explicitly asking for the primary reason for euthanasia may have decreased the need for interviewer interpretation and likely would have improved the agreement between PI-EOLS and EOLS responses.

Despite the multifactorial nature of owners’ euthanasia decisions, the EOLS item addressing reason for euthanasia performed well. Participants appeared satisfied with the response choices provided for this item as <2% chose the response “other, please describe”, and all free-text entries reflected one of the available response choices. Additionally, the EOLS contained two items designed to capture the primary and all additional motivations for pursuing euthanasia. The number of matching responses between PI-EOLS and EOLS was considerably better when all reasons for euthanasia were considered, thus supporting that the EOLS reliably captures all relevant factors in participants’ euthanasia decisions.

Perimortem quality of life is challenging to assess objectively, nevertheless there was moderate agreement between EOLS and PI-EOLS responses for this theme. Unlike the fully open-ended structure of the reason for euthanasia interview question, the perimortem quality of life interview question was open-ended but paralleled the structure of the EOLS by asking participants to describe their dog’s perimortem quality of life “in terms of good days and bad days”. This specific phrasing likely contributed to the higher agreement obtained with this question as compared to the reason for euthanasia question. Among the discrepancies between EOLS and PI-EOLS responses for perimortem quality of life, ten out of eleven were minor, being only one or two degrees off on the 7-point Likert-type scale.

Among the three validation themes, cause of death had the highest kappa statistic representing substantial agreement. Most discrepancies between PI-EOLS and EOLS responses were due to utilization of different survey logic to arrive at the same cause of death or reversal of primary and secondary cause of death between the interview and the instrument. For three remaining discrepant responses, the dogs’ causes of death were vague (undiagnosed illness, undefined cancer, and “old age”) which may have contributed to participant uncertainty and inconsistency in EOLS and PI-EOLS responses. The response choices that the EOLS provides for cause of death appear satisfactory as <5% of participants chose the option “other, please describe” for this item. Furthermore, 75% of these free-text entries reflected responses choices that were provided. Based on the results of this reliability assessment, the EOLS was successful in capturing companion dog cause of death, reason for euthanasia, and perimortem quality of life as described in participants’ proxy interviews concerning their dogs’ deaths.

Review of the free-text responses throughout the EOLS revealed that 80% of participants utilized the option of “other, please describe” for at least one item and/or the option of providing additional narrative information at the conclusion of the survey. Although this represents a significant portion of participants, only 5% of the responses to the twenty “other, please describe” items were provided as free-text. Furthermore, the free-text entries for eighteen of these twenty items did not contain novel, relevant responses that were repeated by multiple participants. This further supports the conclusions of the expert panel and pilot study that the response choices within the EOLS are adequately comprehensive to capture scientifically relevant details surrounding companion dogs’ death. The two items for which the “other, please describe” free-text entries included a frequent, novel, and relevant response were the questions of “Who was present at your dog’s death” and “Who performed the euthanasia”. Based on participant responses, the response choice “another dog” will be added to the question about who was present, and the response choice “a veterinarian or veterinary technician working for an at-home euthanasia or hospice service” will be added to the question about who performed the euthanasia.

Although many participants shared additional narrative information at the end of the EOLS, free-text analysis of these responses did not reveal any themes relevant to the EOLS that were not already addressed elsewhere in the instrument. Further details provided by participants in the additional narrative information either expanded beyond items already contained in the EOLS or broached subjects beyond the scope of the EOLS, the most frequent of which was detailing their personal experience of their dog’s death. Despite this information being outside the scope of the EOLS, we recognize the importance of understanding owners’ perceptions and experiences of their dogs’ deaths. Therefore, additional data collection instruments are under development by the DAP to gather and analyze this information. Additionally, despite the potential for redundancy within free-text responses throughout the EOLS, it is warranted to retain these options in order to maintain participant satisfaction and enable free-text analysis which could guide future EOLS alterations.

This study has limitations. For instance, the EOLS development did not involve interviews with focus groups of participants and/or veterinarians to create items. However, a diverse group of experts collaborated and utilized applicable literature to create the contents of the EOLS, and face validity was evaluated by an expert panel and pilot participants. Additionally, although a post-hoc analysis, the free-text analysis was performed to identify any repeated relevant response choices or themes that were missing from the EOLS. This analysis identified only two items that warranted minor modification.

The purpose of the reliability assessment evaluating cause of death, reason for euthanasia and perimortem quality of life was to determine if participants’ verbal responses during an interview matched their digital responses, which were provided up to 1 month later. Repeatability assessment was not subsequently conducted with the digital version of the EOLS both because of the increasing time lag from the death of the dog, and because discussing an individual’s deceased dog is an innately sensitive topic. To balance the need to demonstrate validity and reliability against the burden of asking participants to repeatedly relive the potentially unpleasant experience of their dog’s death, the authors utilized separate groups of participants for the pilot study and reliability assessment and performed a post-hoc analysis of all participants’ voluntary free-text responses, but did not confirm repeatability of responses to the digital instrument itself.

The length of time between a dog’s death and owner participation in the pilot study or reliability assessment was not standardized and thus the impact of recall bias on participant responses may be variable. However, care was taken to only choose owners who had reported the death of their dog within the month prior to participation and the EOLS requires participants to complete the survey within 40 days of invitation.

Much of the data gathered in the EOLS cannot be independently verified. Consequently, the authors assessed reliability by comparing participants’ verbal and digital EOLS responses for agreement. This qualitative strategy is more subjective than comparative quantitative validation techniques. Additionally, the interview process used to create the PI-EOLS permitted some responses to be broad and unstructured, which required interviewer interpretation of participant’s responses. Despite these challenges, agreement was fair to substantial, and deeper evaluation of discrepancies between PI-EOLS and EOLS responses often revealed only minor differences in survey logic or reversal of primary and secondary choices, indicating that relevant information was captured despite imperfect agreement.

Although survey data are inherently subject to inaccurate recall and misinterpretation,^50^ the information the DAP seeks regarding companion dog death is primarily or exclusively available from dog owners. While the EOLS has proven to be a reliable method for obtaining these data from owners, it is important to interpret these data with awareness of their source. Therefore, for dogs who die with veterinary assistance, the DAP is building processes to collect veterinarian-completed survey data and/or necropsy reports to substantiate and complement information provided by owners.

The EOLS has proven to be a valid and reliable tool for collecting companion dog mortality data and consequently is an appropriate foundation upon which to build the DAP’s companion dog mortality information. We anticipate that the EOLS data will provide new insights into companion dogs’ end-of-life experiences that will empower veterinarians to better care for their terminally ill and geriatric patients.

## Acknowledgments

The authors would like to thank Lisa Moses, VMD, DACVIM, Erica C. Jonlin, PhD, and Amber Keyser, PhD for their participation in the expert panel.

The Dog Aging Project thanks study participants, their dogs, and community veterinarians for their important contributions.

The Dog Aging Project is supported by U19 grant AG057377 from the National Institute on Aging, a part of the National Institutes of Health, and by private donations.

This manuscript is solely the responsibility of the authors and does not necessarily represent the official views of the National Institutes of Health.

The authors declare that there were no conflicts of interest.

## Dog Aging Project Consortium Authors

Joshua M. Akey^1^, Brooke Benton^2^, Elhanan Borenstein^3^, Marta G. Castelhano^4^, Amanda E. Coleman^5^, Kate E. Creevy^6^, Kyle Crowder^7^, Matthew D. Dunbar^8^, Virginia R. Fajt^9^, Annette L. Fitzpatrick^10^, Unity Jeffery^11^, Erica C Jonlin^12^, Matt Kaeberlein^13^, Elinor K. Karlsson^14^, Kathleen F. Kerr^15^, Jonathan M. Levine^16^, Jing Ma^17^, Robyn L McClelland^18^, Daniel E.L. Promislow^19^, Audrey Ruple^20^, Stephen M. Schwartz^21^, Sandi Shrager^22^, Noah Snyder-Mackler^23^, M. Katherine Tolbert^24^, Silvan R. Urfer^25^, Benjamin S. Wilfond^26^

^1^ Lewis-Sigler Institute for Integrative Genomics, Princeton University, Princeton, NJ, USA

^2^ Department of Laboratory Medicine and Pathology, University of Washington School of Medicine, Seattle, WA, USA

^3^ Department of Clinical Microbiology and Immunology, Sackler Faculty of Medicine, Tel Aviv University, Tel Aviv, Israel

^4^ Cornell Veterinary Biobank, College of Veterinary Medicine, Cornell University, Ithaca, NY, USA

^5^ Department of Small Animal Medicine and Surgery, College of Veterinary Medicine, University of Georgia, Athens, GA, USA

^6^ Department of Small Animal Clinical Sciences, Texas A&M University College of Veterinary Medicine & Biomedical Sciences, College Station, TX, USA

^7^ Department of Sociology, University of Washington, Seattle, WA, USA

^8^ Center for Studies in Demography and Ecology, University of Washington, Seattle, WA, USA

^9^ Department of Veterinary Physiology and Pharmacology, Texas A&M University College of Veterinary Medicine & Biomedical Sciences, College Station, TX, USA

^10^ Department of Family Medicine, University of Washington, Seattle, WA, USA

^11^ Department of Veterinary Pathobiology, Texas A&M University College of Veterinary Medicine & Biomedical Sciences, College Station, TX, USA

^12^ Department of Laboratory Medicine and Pathology, University of Washington School of Medicine, Seattle, WA, USA

^13^ Department of Laboratory Medicine and Pathology, University of Washington School of Medicine, Seattle, WA, USA

^14^ Bioinformatics and Integrative Biology, University of Massachusetts Chan Medical School, Worcester, MA, USA

^15^ Department of Biostatistics, University of Washington, Seattle, WA, USA

^16^ Department of Small Animal Clinical Sciences, Texas A&M University College of Veterinary Medicine & Biomedical Sciences, College Station, TX, USA

^17^ Division of Public Health Sciences, Fred Hutchinson Cancer Research Center, Seattle, WA,USA

^18^ Department of Biostatistics, University of Washington, Seattle, WA, USA

^19^ Department of Laboratory Medicine and Pathology, University of Washington School of Medicine, Seattle, WA, USA

^20^ Department of Population Health Sciences, Virginia-Maryland College of Veterinary Medicine, Virginia Tech, Blacksburg, VA, USA

^21^ Epidemiology Program, Fred Hutchinson Cancer Research Center, Seattle, WA, USA

^22^ Collaborative Health Studies Coordinating Center, Department of Biostatistics, University of Washington, Seattle, WA, USA

^23^ School of Life Sciences, Arizona State University, Tempe, AZ, USA

^24^ Department of Small Animal Clinical Sciences, Texas A&M University College of Veterinary Medicine & Biomedical Sciences, College Station, TX, USA

^25^ Department of Laboratory Medicine and Pathology, University of Washington School of Medicine, Seattle, WA, USA

^26^ Treuman Katz Center for Pediatric Bioethics, Seattle Children’s Research Institute, Seattle, WA, USA

